# A Rose by any Other Name: Towards a quantitative theory of memory in neuroimmune memories

**DOI:** 10.1101/2024.07.25.605114

**Authors:** Nabil T. Fadai, Victor Turcanu, Dan V. Nicolau

**Affiliations:** School of Mathematical Sciences, University Park, University of Nottingham, NG7 2RD, UK; Peter Gorer Department of Immunobiology, School of Immunology and Microbial Sciences, King’s College London, Guy’s Hospital, Great Maze Pond, SE1 1UL, UK; Department of Women and Children’s Health, School of Life Course and Population Sciences, King’s College London, St Thomas’ Hospital, Westminster Bridge Road, SE1 7EH, UK

## Abstract

Humanity has known about – and been fascinated by – the connection between mind and body since time immemorial, yet we have only been able to begin to quantitatively explore their interactions in recent years. Even so, the field of neuroimmunology is categorically in its infancy. Despite myriad and diverse experimental reports, no coherent theoretical approach to the neuroimmune system has been even attempted, which makes it difficult to make sense of the emerging body of empirical findings. Here, we take the first steps towards this goal by introducing a mathematical framework that describes the triggering and control of neurological memories (engrams) of immune challenges. Using peanut allergies as a model system, we show how a simple differential equation model of coupled ‘immune-engram’ responses can explain a number of key observations regarding putative neurological control of focal inflammatory responses and failures thereof. Simulations of our model identify four areas of the parameter regime corresponding to distinct consequences of ‘fake’ immune stimulation: a) resolution b) finite oscillations of inflammation, followed by resolution c) sustained, self-amplifying oscillations (cytokine storm-like phenomena) and d) resolution followed by the permanent establishment of a chronic higher-baseline inflammatory state. Importantly, we then show how our model recapitulates the key qualitative features of a recent experimental description of immune engram encoding and re-activation (respectively, suppression) in a mouse model of colitis. We conclude with remarks around clinical implications as well as directions for future work in theoretical neuroimmunology.

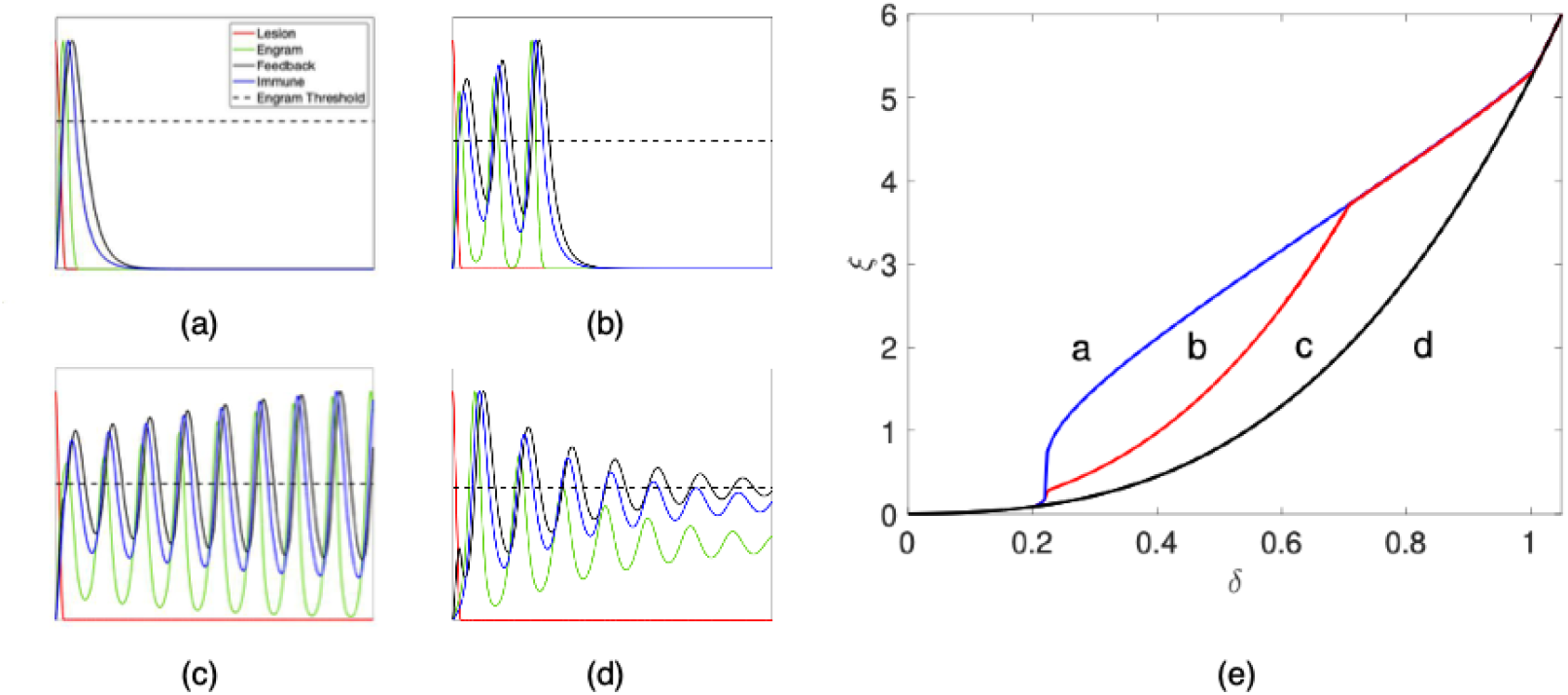

## Introduction

Following the emergence of metazoa some 600m years ago (Tikhonenkov et al., 2020), multicellular animals rapidly increased in complexity and consequently needed to process increasingly complex information about their environments and communities in order to survive. Molecular and cellular mechanisms for sensing, integrating and appropriately responding to environmental challenges evolved concurrently in both the nervous and immune systems (Ottaviani et al., 2007). A growing body of evidence now indicates that shared evolutionary imperatives in fact led these systems to co-evolve to protect the organism from pathogens and other dangers as well as to implement homeostatic control (Kraus et al., 2021). Cooperation between the nervous and immune systems is now known to take place at a number of levels and in each case bidirectionally (Kamimura et al., 2020). For example, neuronal signals optimise the peripheral immune response (Jin et al., 2024), while traditionally “immune” processes are increasingly recognised to play indispensable roles in the growth, development and maintenance of neural tissues and the brain itself (Gruol, 2023).

Tellingly, while these systems have historically been studied separately both scientifically and clinically, the list of molecules with dual functions in both systems and in both health and disease is long and growing (Eberl, 2022). Cooperation and coordination between the immune system and the brain has, indeed, been described for millennia – long before the former was properly identified as a separate entity. The idea that health of one is not possible without the other is captured, for example, in the Latin maxim “*mens sana in corpore sana”*. Formal clinical and scientific descriptions of neuroimmune interactions have historically been largely anecdotal, however. Writing in 1886, Mackensie described a case of severe, clinically objective hayfever triggered by an artificial rose (Mackensie, 1886). In the 20th century, Ader and Cohen described behaviourally conditioned immune suppression, so severe as to cause the death of a number of the animals involved (Ader and Cohen, 1975). These and myriad other experimental findings have remained isolated from one another and no theoretical description of the neuroimmune system has even been attempted.

On the other hand, recent technological advances have, for the first time, allowed more detailed and formal experimental investigations of neuroimmune interactions in animal models. Optogenetic methods have, recently, for example, provided strong evidence that immune ‘engrams’ – neuronal memories of earlier peripheral inflammation responses – are potentially stored following a disease state and can be both reactivated and suppressed directly through stimulation of the originally activated population of neurons (Koren et al., 2021). Progress over the last few years has specifically identified a number of brain areas involved in neuroimmune process, particularly pointing to the insular cortex as being a key area for the integration, storage and activation of peripheral immune responses as well as for simulation of potential future physiologic states (Livneh et al., 2020). However, justifying the key mechanistic behaviour of engram activation, as well as the immune system’s response, has been largely unexplored.

Here, we take the first steps towards a coherent theoretical description of neuroimmune memories or “engrams”. We briefly review evolutionary studies on the co-evolution of the neural and immune systems as well as summarising recent experimental evidence regarding neuroimmunity in peripheral organs, such as the gut. Next, we present a plausible, general mathematical framework for studying neuroimmune engrams, showing how this framework can capture, explain and reconcile – at a high level – some of the key relevant experimental findings. Finally, we discuss clinical implications as well as prospective methods for therapeutically modulating the neuroimmune system; and suggest future directions of work for this nascent field of study.

### The Evolutionary History of Neuroimmune Interactions

The evolutionary role of the brain for sensing and acting on danger precedes the emergence of an organised immune system and certainly of the adaptive immune system, which first appeared in jawed fish, some 500 mya (Flajnik and Kasahara, 2010). The defensive role of the brain, leading to the formation of engrams in higher organisms, has probably emerged under evolutionary pressures from microbes, bringing together coordinated responses from pattern recognition receptors, soluble factors and ion channels that evolved over hundreds of millions of years (Kraus, 2021). The subsequent development of the specialised immune system, which identifies and responds to non-self antigens, came about in close coordination with pre-existing neuron-performed defences.

Thus, even primitive organisms make full use of neural-coordinated responses for defence. *C elegans* uses a behavioural reaction as the first line of defence for evading pathogenic bacteria (Schulenburg, 2004). *C elegans* has only limited immune defences as it has some conserved immune signalling pathways but lacks inducible nitric oxide synthase (Wibisono and Sun, 2021). This is presumably why neurons take an active role in anti-microbial defence. Cao et al (2017) distinguished two groups of *C elegans* neurons that express OCTR-1, a neuronal G-protein-coupled receptor analogous to the human norepinephrine receptor.

One neuronal group controls innate immune pathways involved in microbial killing, while the other group promotes pathogen avoidance behaviour (Cao et al, 2017). Octopamine, dopamine, serotonin, and other neurotransmitters provide aminergic signalling from neurons to innate immunity effectors in insects like *Drosophila* (Cattabriga, 2023) or the beet armyworm *Spodoptera* (Kim, 2010). They ensure the integration of neuronal responses with immune effectors such as the circulating hemocytes that can phagocyte bacteria and form defensive nodules. The *Drosophila* brain also maintains a direct role in defence, so that, for instance, a conserved olfactory circuit dedicated for detecting harmful microbes was observed (Stensmyr, 2012).

The immune system subsequently underwent significant evolutionary developments, so that fish display strong protective mucosal immunity responses. The nasal immune structures organised as the nasopharynx-associated lymphoid tissue (NALT) in bony fish are closely associated with the olfactory organ that is connected to the brain via the olfactory bulb (Kumar Das, 2020). Teleost NALT responds to bacterial infections of the olfactory organ and also to nasal vaccinations by mounting a humoral response relying upon the nasal IgM and IgT B cell repertoires. The close proximity between the NALT immune cells and olfactory neurons leads to neuron responses after vaccination, as shown in rainbow trout – thus, crypt neurons from the olfactory organ enter apoptosis rapidly after vaccination but recover within a few days (Sepahi, 2019).

Strong integration between brain effectors and the immune system is further enhanced in all vertebrates by neural control of corticosteroid secretion through the hypothalamic-pituitary-adrenal axis and the autonomic nervous system which can suppress excessive or chronic immune responses (Nicolaides, 2015). More recently, Koren and colleagues demonstrated, in a mouse model, that insular cortex neurons encode and retrieve specific responses in the gut, and that their re-activation can reproduce an autoimmune-type immune reaction even in the absence of any toxic challenge (Koren, 2021).

### Neuroimmunological engrams drive allergic and autoimmune reactions

Neuroimmune phenomena are extraordinarily diverse, involving virtually every major chronic disease state, including pain (Meade and Garvey, 2022), cancer (Scheff and Saloman, 2021), mental illnesses (Hodes et al., 2015) and cardiovascular disease (Zarate et al., 2024). In order to take concrete steps towards the development of a theoretical framework for describing neuroimmune interactions generally, we focus here – partially in a nod to the seminal historical work of Mackensie – on a (relatively) simple model system: peanut allergies.

Peanut allergy is one of the most common food allergies in adults. Its overall self-reported lifetime prevalence in Europe is 1.5% of the population (1.0-2.1 in various studies) but its point prevalence when verified by a peanut challenge is much lower at 0.1% (Spolidoro et al., 2023). Peanut allergy lifelong persistence being around 80% of patients, its potential to cause severe or even fatal anaphylactic reactions and the limited success of current immunotherapies with providing a definitive cure, together, make it a significant public health concern (Riggioni, 2024). Nevertheless, a prospective randomised controlled trials of early introduction of peanut products in babies’ diets to induce oral immune tolerance led to over 80% peanut allergy prevention in highly atopic children (Du Toit, 2015 and Du Toit, 2016). These studies consequently led to the changes of worldwide guidelines, replacing the advice to avoid peanuts until 3 years of age with early-life peanut consumption (Du Toit, 2015 and Turcanu, 2017). Therefore, peanut allergy may decrease in future generations, whilst still manifesting in older children and adults who had not been protected through early-life peanut consumption.

Indeed, peanut-allergic individuals flying on airplanes occasionally report allergic reactions, sometimes requiring adrenalin treatment, usually starting after they perceive the smell of roasted peanuts eaten by other passengers (Sicherer, 1999 and Greenhawt, 2009). These reactions appear paradoxical, as peanut antigens are almost undetectable in the air because peanuts are ‘sticky’ and lipid-rich. Indeed, aerosolised peanut antigens were undetectable even in a study where participants wore air monitors on their heads whilst opening up to 15 bags of peanuts and consuming the contents, in order to simulate the environment of a commercial airline (Perry, 2004). It has also been found that airborne peanut allergens are below the detection level of a very sensitive ELISA assay. The only situation in which a (very weak) signal could be detected was when air was sampled directly above peanuts whilst these were being deshelled (Brough, 2013).

More recently, it has been reported that airborne peanut allergens are below the detection threshold, when 84 peanut allergic children were placed in a small room only 0.5m in front of a peanut bowl containing 300g of roasted peanuts. None had a severe or moderate allergic reaction and only two had minor rhino-conjunctivitis (Björkman, 2021). Likewise, no reaction was observed when peanut allergic individuals underwent inhalation challenges where the smell of peanut was masked. In these challenges, peanut butter was placed in a gauze-covered box alongside tuna and mint to mask odours and the box was kept 12 inches from the patient’s nose for 10 minutes (Simonte, 2003). Importantly, perceiving the smell of peanuts does not necessarily entail exposure to peanut allergens because the odour of roasted peanuts is not caused by peanut allergens (which are relatively large, kilodalton-sized proteins) but by a group of more than 40 small-molecule, volatile compounds. These are mainly pyrazines, pyrrolines, phenols and butanoic acid derivatives whose molecules are simply too small to cross-link specific IgE antibodies on the surface of mast cells or basophils and trigger histamine release (Kaneko, 2013). Taken together, these and other studies suggest that the allergic reactions reported by peanut allergic patients who perceive the odour of peanuts cannot be triggered through the cross-linking of specific IgE antibodies by peanut allergens and may, therefore, be largely attributable to a – hitherto putative – immune engram created and reinforced by allergic reactions from the past.

An analogous situation was reported almost 140 years ago, when Mackenzie described the induction of an allergic reaction, which he termed ‘rose cold’, by showing an artificial rose to a patient (Mackenzie, 1885). In this case, Mackenzie obtained an “artificial rose of such exquisite workmanship that it presented a perfect counterfeit of the original. To exclude every possible error, each leaf was carefully wiped, so that not a single particle of foreign matter was secreted within the convolutions of the artificial flower.” Briefly, minutes after seeing this artificial flower, the patient experienced tickling and intense itching in the back of the throat, red and runny nose and eyes and finally a feeling of ‘oppression in the chest’ with ‘slight embarrassment of respiration’. Paradoxically, this allergic reaction was not caused by the exposure to an actual allergen, but merely by the visual perception of its presence which then triggered the corresponding pathogenic engram which subsequently reproduced an allergic reaction.

Further evidence of pathogenic engrams driving allergic and autoimmune reactions emerged since this report. This has led to a wider belief that when neuronal networks sense danger, they build up neurological memories that may reproduce complex neuroimmune responses upon restimulation. Such neurological ‘engrams’ would have clear roles in the defence of the organism, for example by accelerating the initial immune response to pathogens and other dangers. Cases such as those described above, however, additionally convincingly demonstrate that immune engram restimulation by spurious signals such as peanut small molecule smells or seeing an artificial rose, may trigger pathogenic responses that are potentially indistinguishable from the ‘native’ peripheral immune response itself.

Therefore, it seems reasonable to hypothesise that these reactions likely result from the activation of neuroimmunological memories (‘engrams’) that were formed when smelling the odour of peanuts was repeatedly associated with allergic reactions caused by accidental peanut ingestion (Figure 1A). Once established, activation of these engrams through the smell of peanuts (in the absence of an actual exposure to peanut allergens) initiates neural-driven pathogenic responses that reproduce allergic reactions, following a clinical course similar to reactions having occurred in the past following genuine ingestion of peanut antigens. These interactions would be co-mediated with the peripheral immune response, forming a network of interactions illustrated in cartoon format in Figure 1B. We are here using peanut allergies as a fairly well circumscribed model system, but the reader will appreciate that the processes we are investigation would occur in a very wide range of situation in which an immune response is triggered.

**Figure 1:**
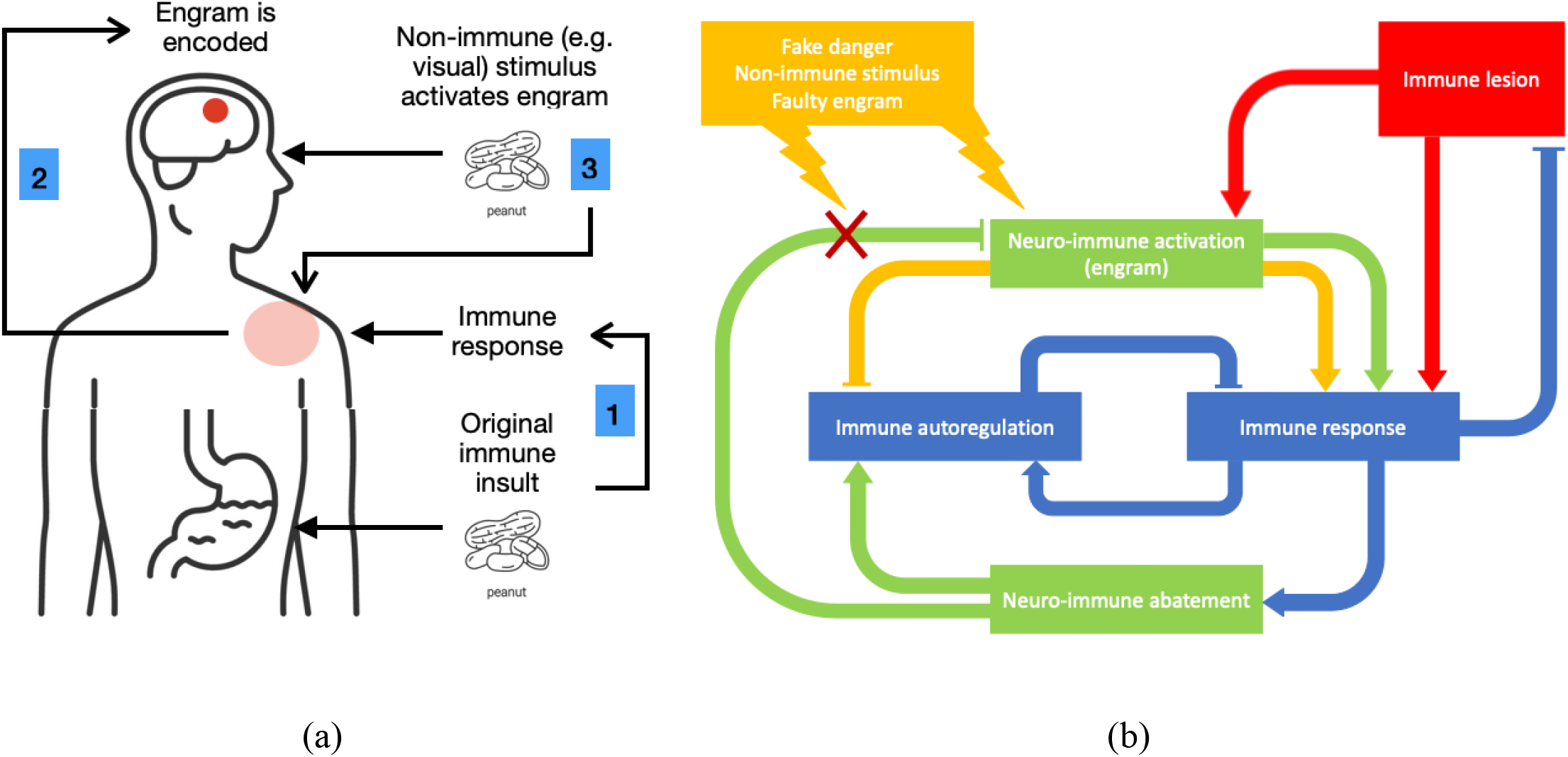
**(a)** A simplified model of the potential engram-inflammation feedback network. An external immune lesion causes an inflammatory response as well as activation of a neural engram, which can be reactivated by either direct or indirect stimulation. **(b)** Typical inflammation responses would resolve the immune lesion, followed by an abatement of immune and engram networks via feedback mechanisms. However, in the case of faulty engrams, or other engram-activating stimulation, the feed-back mechanisms are disrupted, causing a combination of immune hyperactivation and/or damage to immune autoregulation.

### Mathematical Model of Neuroimmune Engram Effects on Peripheral Immune Responses

We now propose a differential equation model for the mechanism that underlies the role of a pathogenic engram, reproducing the symptoms of an allergic or autoimmune reaction. The outlines of our model are captured in Panel B of Figure 1. We will refer to this ODE system as the ‘Engram-Immune Model’ (EIM) and will examine particular qualitative features arising from this model below.

Figure 2 depicts the proposed paradigm in which a clinically manifest allergic reaction (for example to peanuts in an allergic patient) results from the contribution of the brain-based engram effect added up to the body-based immune response to peanut allergens. Allergic reactions are defence responses designed to protect the organism against allergens that are misperceived as similar to dangerous toxins, viruses, bacteria etc. Thus, in our model, the neuronal network storing the engram encodes the memory of past allergic reactions to peanuts together with memories of olfactive, visual, taste flavours etc. associated with the presence of the peanut allergens. When one or more of these associated signals activate the neuron network, the engram effect brings out the memory of previous allergic reactions, perhaps in order to prime the organism for the purpose of providing a rapid defensive response, if and when peanut allergens are actually present and trigger the degranulation of tissue mast cells and blood basophils with subsequent release of immune mediators such as histamine, serotonin and various cytokines and chemokines.

**Figure 2:**
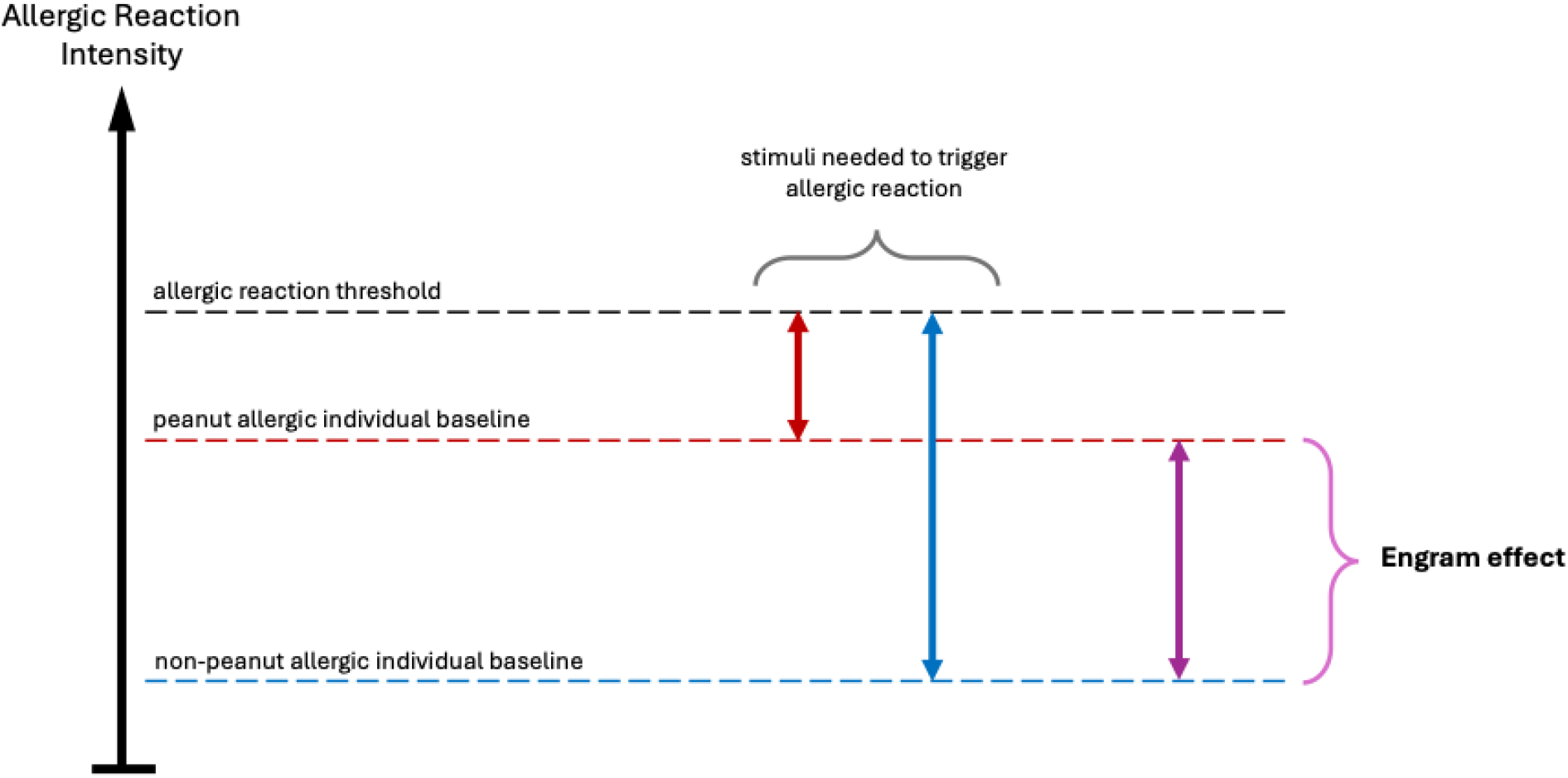
A simplified model of the contribution of the engram-encoded effect in triggering allergic reactions. Typically, the neuron network retains as an engram the memory of previous allergic reactions and its association with olfactive, visual, taste flavours etc signals associated with the presence of the allergen. When activated by such non-allergenic stimuli, the neuronal network brings the body to an engram-primed sub-clinical reactive status. Thus, the peripheral immune effectors are being primed for a subsequent full-blown clinical allergic reaction if exposed to even minimal levels of the offending allergenic stimulus, for the purpose of providing a rapid, evolutionarily-beneficial defensive response.

The EIM combines various biophysical mechanisms that interact with each other through engrams, the immune system, and external stressors. To couple interactions of putative immune engrams, induced either by physiological (peripheral) or central challenges, i.e. both “real” and “indirect”, with responses from the immune system, we consider a simplified four-compartment mathematical model whose concentrations evolve in time alone: the engram-promoting signal *x*, which promotes an immune response, an engram-inhibiting signal *z*, which inhibits *x*, the immune/pro-inflammatory response *y*, and an immune lesion *u* which serves as the catalyst to the arising dynamics. Here, *u* can be viewed as some external stimulus, such as a virus or an allergen. A summary of the mechanistic responses is shown in Figure 1B; further details of mechanistic model choices can be found in the Supplementary Information.

Under suitable non-dimensionalisation of variables and mechanistic processes, the EIM can be described via a system of four ordinary differential equations (ODEs):

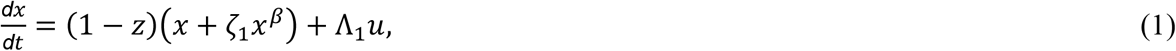

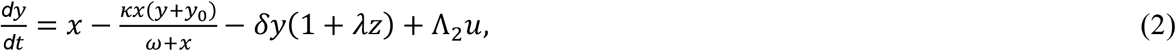

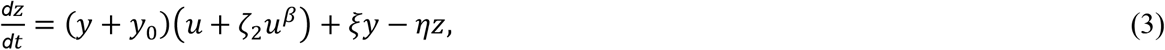

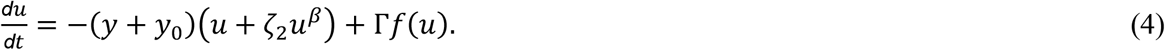

The engram-promoting signal, *x*, activates below a threshold value of the inhibition signal *z*, as well as in the presence of the lesion *u*. The immune response *y*, in turn, is activated by the presence of the engram, but is hindered by a combination of autoimmune disregulation and/or natural decay from the engram-immune feedback. The engram-inhibiting signal, *z*, is activated in the presence of elevated immune activity, as well as resolution of the immune lesion. The immune lesion leads to a response by the immune response, but can self-replicate in instances of viral infection (e.g., COVID-19). In the case of allergic responses (to a single stimulus), this self-replication term does not appear in the equations.

A crucial component of the mathematical modelling is the ‘cut-off’ mechanism appearing in the engram and lesion equations. In particular, the inclusion of the terms *ζ*_1_*x*^*β*^ and *ζ*_2_*u*^*β*^ allow for smooth transitions to their zero-states (the absence of an engram/immune lesion), for 0 < *β* < 1 and particular choices of *ζ*_1,2_. This cut-off mechanism extends the notion of engram activity being proportional to the concentration of population present, but prevents the existence of an exponentially-small population that suddenly resurges later in time. In population dynamics, this is sometimes referred to as the ‘atto-fox problem’ (Murray et al. 1986; Mollison, 1991; Lobry and Sari, 2015), in which an infeasibly small population persists and can resurge to its original population size; an equivalent ‘yocto-cell problem’ (Fowler, 2021) has been coined for mathematical models at the cellular scale. Similar cut-off mechanisms could be incorporated for the immune system and inhibition signal, but are omitted for brevity and simplicity. We will refer to this ODE system, coupled with the initial condition (*x*(0), *y*(0), *z*(0), *u*(0)) = (0,0,0, *V*_0_) as the ‘Engram-Immune Model’ (EIM), and we will examine particular qualitative features arising from this model below.

### Qualitative Features of the EIM

As mentioned previously, the EIM combines various mechanisms that interact between engrams, the immune system, and external stressors. Naturally, it is important to understand the qualitative features that arise in various parameter regimes of the EIM. To examine these features, we numerically compute solutions of the EIM using MATLAB’s ode15s function and scale all variables (*x, y, z, u*) relative to their maximum value. This rescaling of variable height is particularly plausible in the absence of further experimentally-determined parameter values; with reference to the Supplementary Information, each variable in the (dimensionless) EIM can be rescaled back to dimensional values independently of other variable scalings.

In Figure 3, we observe four main qualitative features of the EIM, with each parameter set corresponding to a different fictitious patient.

**Figure 3:**
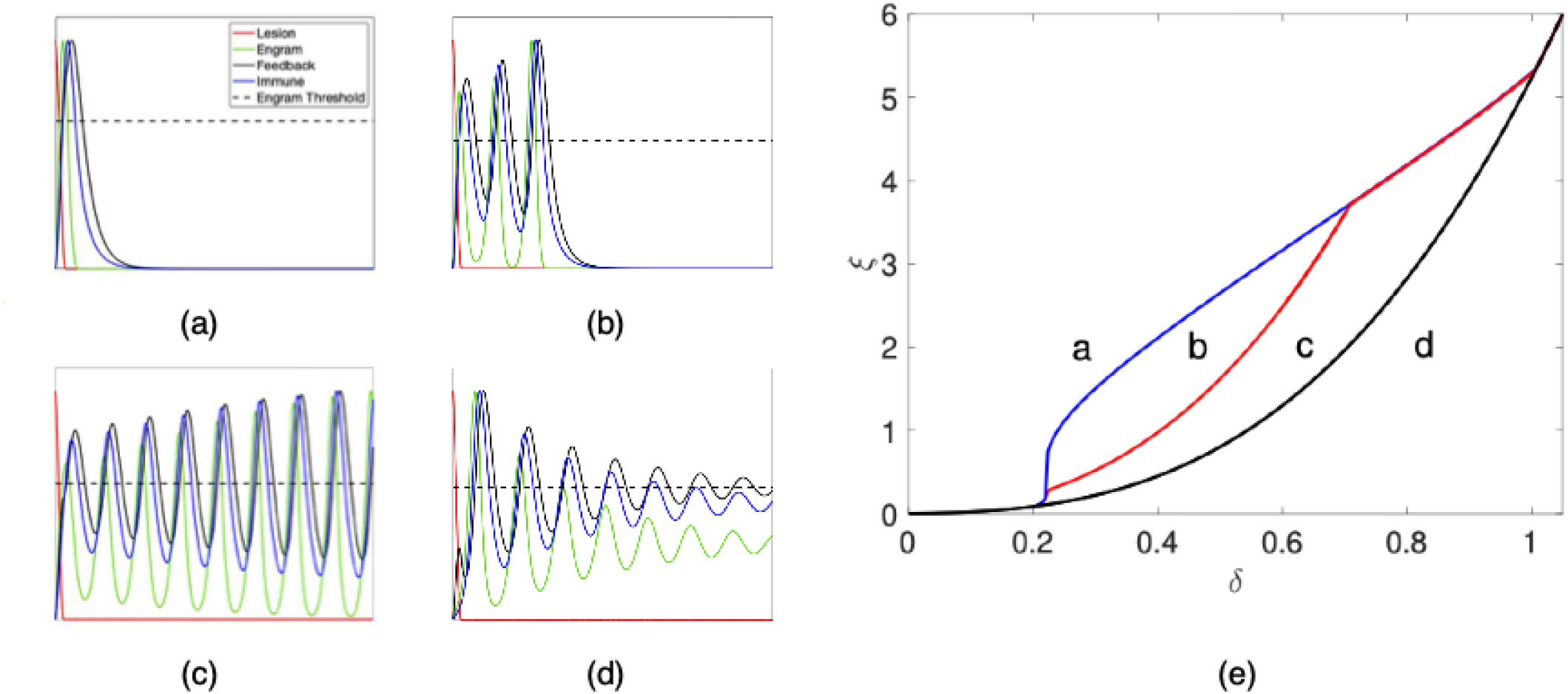
Qualitative features and transitions of the EIM. Simulations correspond to qualitative neuro-immune responses as functions of time (arbitrary units), in the presence of an immune lesion (red). All quantities shown are rescaled relative to their maximum quantities and all parameters are listed in Table 1. In a healthy patient response (a), a single spike of engram-promoting activity (x, green) gives rise to elevated immune activity (y, blue) and engram-inhibiting activity (z, black). Inhibition signal activity above the engram threshold (dashed black) reduces engram-promoting activity, which in turn resolves immune and inhibition activity. For slower inhibition responses (smaller ξ), the engram-promoting activity is prematurely reduced, causing multiple spikes of immune activity (b) or a sustained pattern of spiking behaviour (c). For very strong feedback activity, a sustained state of elevated inflammation is created (d). The bifurcation diagram (e) shows the transitions between EIM phenotypes, relative to the engram threshold (characterised via the proxy variable ξ) and the immune response’s clearance rate (characterised via the proxy variable δ). The boundary of sustained inflammation (black) corresponds to the Hopf bifurcation described in Equation (7). The ‘terminating spike criteria’, described in Equation (11), determines the single-to-multiple spike transition (blue) and the boundary of infinite spiking (red).

1. In the parameter set corresponding to healthy patient response, the engram resolves the immune lesion completely and the patient experiences a single ‘spike’ of heightened immune activity. Instances of this parameter regime would also correspond to a mild viral infection or a localised allergic sensitivity.
2. However, for parameter sets corresponding to faulty engrams, the engram-immune network becomes imbalanced in its positive-negative feedback loops.
3. In moderate cases of network imbalance, a patient may experience several spikes of heightened immune activity, followed by a complete resolution of the immune lesion. Noting that the EIM timescale is also non-dimensional (in arbitrary time units), the frequency of these spikes could vary, clinically or sub-clinically, from the order of hours to years, depending on the particular affliction.
4. Finally, in severe network disruption, including a severe allergic reaction or an auto-immune disorder, the patient experiences sustained, self-amplifying oscillations (cytokine storm-like phenomena), or a permanent establishment of a chronic, elevated inflammatory baseline. Given the wide range of qualitative features that can arise from the EIM, we conclude that this simplistic modelling framework provides a convincing template for describing the engram-immune interactions within various patient cohorts.

**Table 1:** Summary of (dimensionless) variables and parameters used in the Engram-Immune Model. The values of ξ correspond to simulations appearing in the four subfigures of Figure 3.

| Variable / Parameter | Biological Interpretation | Variable / Parameter | Biological interpretation |
| --- | --- | --- | --- |
| $x(t)$ | Engram-promoting signal strength | $\delta = 0.4$ | Inflammation clearance rate |
| $y(t)$ | Immune system response strength | $\lambda = 0.2$ | Inhibition-promoted inflammation clearance rate |
| $z(t)$ | Engram-inhibiting signal strength | $y_0 = 0.2$ | Baseline immune system activity |
| $u(t)$ | Immune lesion strength | $\xi = \{2.2, 1.5, 0.8, 0.1\}$ | Inhibition promotion from inflammatory response |
| $\zeta_1 = 0.75$ | Engram cut-off strength | $\eta = 1$ | Inhibition signal autoregulation |
| $\zeta_2 = 1$ | Lesion cut-off strength | $\kappa = 0$ | Autoimmune damage strength |
| $\beta = 0.5$ | Cut-off strength power law (engram and lesion) | $\alpha = 1$ | Autoimmune damage strength gradient |
| $\Lambda_1 = 0.5$ | Engram promotion from immune lesion | $\Gamma = 0$ | Immune lesion growth rate |
| $\Lambda_2 = 0.5$ | Inflammation promotion from immune lesion | $f(u)$ | Immune lesion growth function |

### Quantifying the transitions of spiking activity in the Engram-Immune Model

As seen in Figure 3, the qualitative features of the EIM naturally divide into four separate response phenotypes. Given the multitude of parameter appearing in the EIM, it is natural to ask what parameter combinations give rise to each kind of response phenotype. However, due to cut-off mechanism appearing in the EIM, local analysis when the engram is nearly zero fails. Therefore, we need to explore different criteria to establish when the cut-off mechanism drives the engram to zero.

To begin, we use standard local analysis about the non-zero steady-state value (*x*^*^, *y*^*^, *z*^*^) in the absence of any immune lesion, i.e. *u* = 0. This steady-state corresponds to the sustained inflammation phenotype described above. For simplicity, we will exclude any auto-immune damage effects from the steady-state calculations here, i.e., *κ* = 0, and obtain the following steady-state when all temporal derivatives are set to zero:

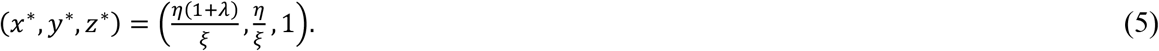

Linearising the EIM about this steady-state provides the Jacobian matrix

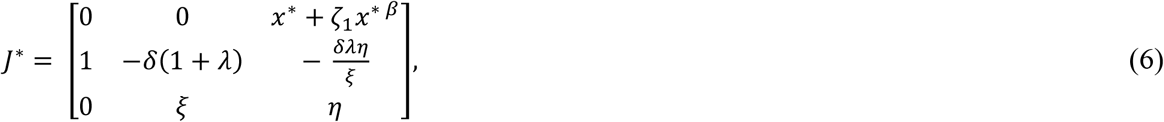

and a Hopf bifurcation (signifying the threshold of this steady-state’s stability) occurs when |*J*^*^ − *iωI*| = 0, where *ω* represents the sustained spiking frequency and *I* is the identity matrix. Solving this complex-valued equation provides the following stability criterion of the sustained inflammation phenotype:

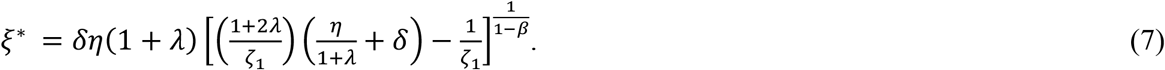

In other words, a critical value *ξ*^*^ separates parameter space into a region where fluctuations of inflammation become a chronic state of elevated inflammation, and a region where spikes of inflammation occur.

As seen in Figure 3, it is possible to have a single spike of inflammation, multiple (but finite) number of inflammation spikes, or a state of continuous spiking activity. To quantify the distinctions between these three phenotypes, it is essential to examine the behaviour of the engram activity *x* in the presence of the elevated inhibition signal, *z*. In the absence of immune lesions, Equation (1) reduces to

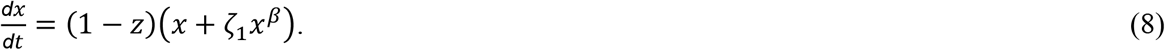

In particular, we note that a single spike of activity includes a time period whereby *z*(*t*) > 1, and thus *x*^*′*^(*t*) < 0. Therefore, we consider the sequence of time intervals *T*_*i*_, in which the inhibition signal is above its activation threshold:

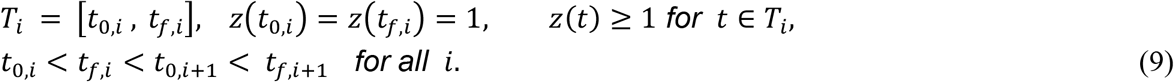

Noting that the cut-off function in *x* allows the engram signal to smoothly transition to zero, we have that each time interval *T*_*i*_ has the potential to have *x*(*t*) ≡ 0 for *t* > *t*_*f*,*i*_, allowing both *y*(*t*) and *z*(*t*) to decay to zero and terminate the spiking behaviour. With this in mind, we integrate Equation (8) from *t*_0,*i*_ to *t*_0,*f*_ and impose the criterion that *x*(*t*_*f*,*i*_) = 0 to establish the end of the spiking activity:

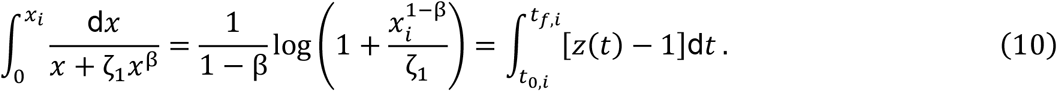

Here, we denote *x*_*i*_ = *x*(*t*_0,*i*_) for simplicity. Finally, we note that the integral in *t* represents the maximum time interval for *x*(*t*) to become identically zero, and thus we are able to determine when the final spike occurs:

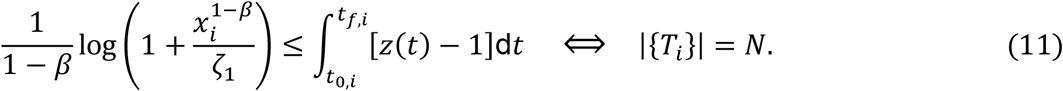

In other words, the first time interval *T*_*N*_ at which Equation (11) occurs tells us that exactly *N* spikes of engram activity occur before *x*(*t*) ≡ 0. While computing the time intervals *T*_*i*_ and the *z*(*t*) integral are both analytically infeasible, they are both straightforward to numerically compute by means of computational ODE solvers in Matlab.

Knowing the ‘terminating spike criteria’, shown in Equation (11), we are also able to determine the region of parameter space where a single spike of engram activity (*N* = 1) transitions to multiple or infinite spiking activity (*N* > 1 or *N* = ∞). As we see in Figure 3, the ‘single spike’ phenotype can transition into the ‘multiple spike’ phenotype or directly into the ‘infinite spiking’ phenotype, depending on parameter values. Furthermore, the ‘single spike’ phenotype can directly transition into the sustained inflammation phenotype, since the multiple-spike and infinite-spike regions only exist for specific choices of parameter values. While the bifurcation diagram in Figure 3 is a small subsection of different parameter choices, it nevertheless highlights the variety of phenotypes within the EIM.

### Incorporating autoimmune damage in the EIM

We conclude this mathematical model section by briefly mentioning the effects of incorporating autoimmune damage in the EIM by setting *κ* > 0. The non-zero steady-states (*x*^*^, *y*^*^, *z*^*^) will have a different equilibrium engram level of *x*^*^, which solves the nonlinear equation

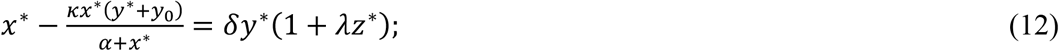

however, the values 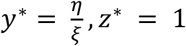 remain unchanged. The arising Jacobian analysis, and hence the Hopf bifurcation criteria, is largely the same, though it incorporates much more tedious algebra. Furthermore, a similar transition between EIM phenotypes to Figure 3 is observed as *κ* is increased. Therefore, we conclude that an increase in the effect of autoimmune damage caused from the engram feedback loop can cause a patient to sustain more inflammatory spikes, or even transition to a chronic inflammatory state.

### The EIM Reproduces Qualitative Experimental Results on Immune Engrams

Next, we sought to use the EIM to model the experiments of Koren et al. (2021). Briefly, a mouse model of colitis was used in this study. Using activity-dependent cell labelling, (FosTRAP), neuronal ensembles in the insular cortices of the animals active under two different inflammatory conditions (dextran sulfate sodium [DSS]-induced colitis and zymosan-induced peritonitis) were identified. Following clinical recovery, chemogenetic reactivation of these neuronal ensembles was sufficient to broadly retrieve the inflammatory state under which these neurons were captured. Furthermore, and fascinatingly, experimentally induced reduction in InsCtx activity (during treatment with DSS) significantly reduced clinical symptoms, such as colon length, spleen weight and many (but not all) histological and inflammatory marker parameters.

To explore the potential of the EIM to capture some of these phenomena, we performed computer experiments of the three experimental regimes described by Koren and colleagues: a) stimulation of an ‘engram-naive’ system with a peripheral inflammatory challenge (such as DSS), b) reactivation of an engram absent a focal insult and c) suppression of engram activity while stimulating with the peripheral immune challenge. The results are shown in Figure 4, labelled “Peripheral Insult”, “Activation” and “Suppression”, respectively. The Peripheral Insult, i.e. naive stimulation experiment shows, as expected, an immune activity spike in response to an immune challenge. The “Activation” experiment, which assumes a stable neuroimmune engram, recapitulates the reactivation experiment of Koren et al.: absent any immune lesion input (*u*, in our model), the total engram activity (*x+z*, in our model) is forced above the engram threshold and leads to an immune response (*y*, in our model). Lastly, in the “Suppression” experiment, we clamped the total engram activity term to zero while providing an immune insult – remarkably, this leads to a reduced (but not absent) immune response, qualitatively similar to the experimental description.

**Figure 4:**
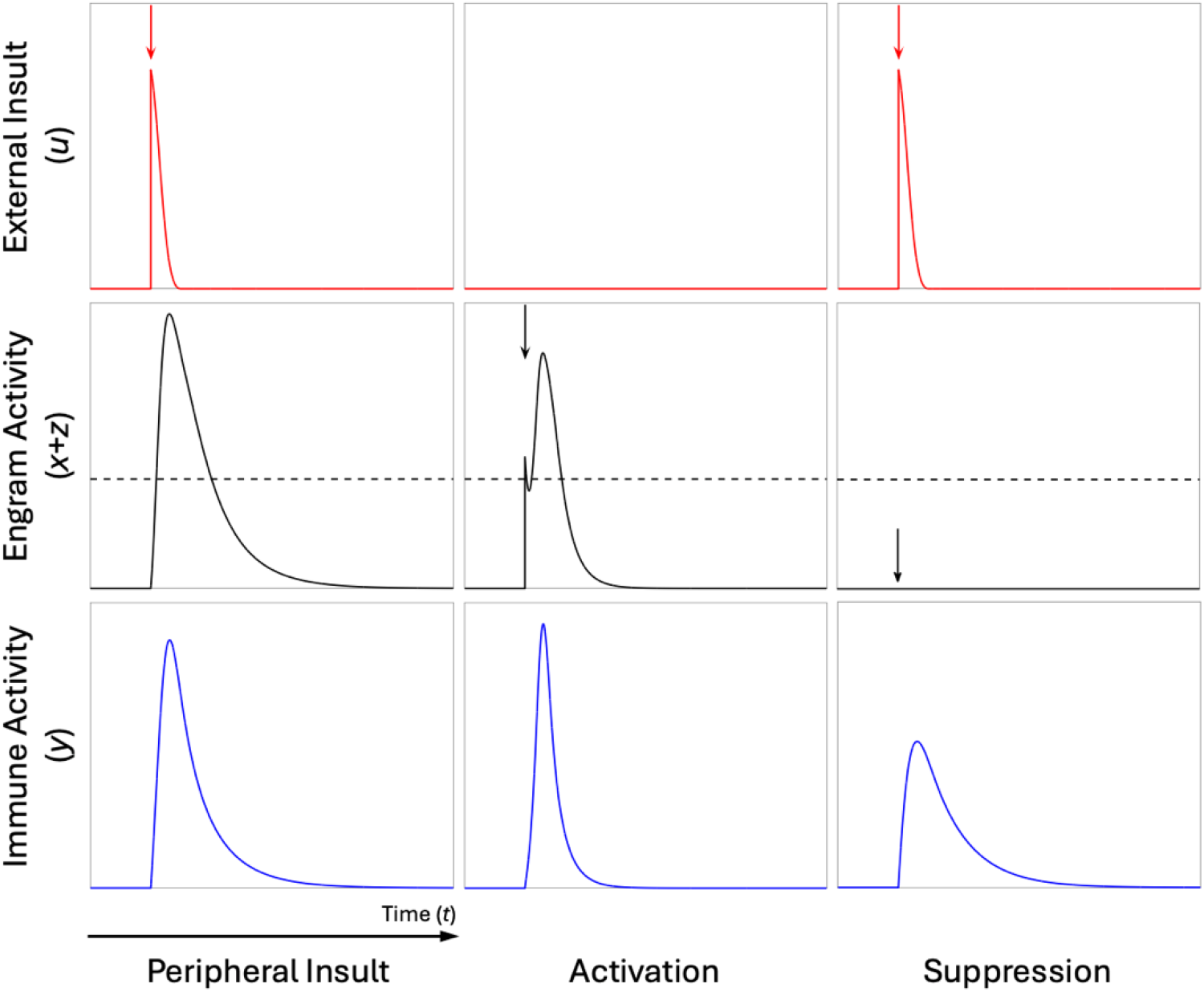
Model simulation of ‘immune engram’ activity recapitulates the experimental results of Koren et al. (2021). Left panel (Peripheral Insult): A peripheral immune challenge (red spike) causes engram activity (black curve) as well as a peripheral immune response (blue curve). Centre panel (Activation): Absent any peripheral challenge, direct stimulation of the engram term in the model (black spike and subsequent activity) above the engram threshold causes a peripheral immune response (blue curve). Right panel (Suppression): In the presence of an immune challenge (red spike) but clamping to zero of engram activity in the model (black flat line, below the engram threshold), leads to a substantially reduced peripheral immune response (blue curve). Parameter values of all experiments shown correspond to the ‘single spike’ parameter set used previously (see Table 1). The engram threshold is shown as a dashed black line and is relative to the total engram activity (solid black line; x+z in the EIM).

## Discussion

Since the brain and immune systems (not necessarily in that order) are the most complex systems known to humankind, development of a theory describing their cross-interaction is clearly a mammoth challenge. Nonetheless, as experimental evidence accumulates from diverse sources and given the clinical importance of understanding neuroimmunity, there is an imperative need for theoretical approaches to the neuroimmune system. These efforts can, *inter alia*, inform the design of potentially fruitful experiments as well as organise interdisciplinary initiatives, including in the clinical setting.

Here, we provide a new mechanistic mathematical model system that provides key qualitative descriptions of the interplay between (still putative) neuroimmune engrams, the immune response, and peripheral immune lesions, such as allergic reactions or viral infections. Crucially, in our model, a ‘cut-off’ mechanism in the immune lesion and engram dynamics allow the system to resolve on its own in finite time; while largely theoretical, the spiking behaviour observed in the immune response is biophysically realistic. Additionally, we observe parameter regimes of this Engram-Immune Model (EIM) that give rise to continuous spiking behaviour in the immune system, in spite of the immune lesion no longer being present. This emergence of self-amplifying spiking behaviour (‘cytokine storm’-like phenomena) and a persistent elevated state of inflammation (observed in autoimmune disorders) supply biological realism and breadth of applicability of the EIM to various settings.

Interrogation of our mathematical model suggests, in the context of allergy, that a clinically manifest allergic reaction is the sum of the effects of (i) allergen-triggered histamine and other mediator release in the body plus (ii) the engram-driven amplifying (or attenuating) effects which are exerted through the vagus nerve, potentially by corticosteroid release or other neuronal effectors that act on the cellular immune effectors in tissues. More generally and importantly, we were also able to use the EIM to qualitatively recapitulate a detailed recent experimental description of immune engram existence, reactivation and suppression in the context of colitis. Further exploration of the capabilities of the EIM to make specific predictions on diverse neuroimmune memory-driven phenomena will form the subject of future work. However, we wish to stress here that this simulation result strongly suggests that a class of models of the EIM type is likely to be broadly applicable to myriad clinical phenomena.

Indeed, putting aside the exciting possibilites of using models of the type described here to build detailed, mechanistic models of neuroimmune interactions – perhaps building towards a general theory – there are also a number of long-term clinical implications from these early results. For example, in the context of peanut allergy, when patients perceive the early-warning symptoms of a developing allergic reaction, it may be possible to act on the engram (via transcranial magnetic stimulation, for example) to stop or ameliorate the reaction. For example, an asthmatic who feels early warning signals that an attack is coming (because of cough episodes, slight tiredness, being nervous and irritable) may use transcranial stimulation to disrupt the engram in addition to using the salbutamol broncho-dilating inhaler. Likewise, in addition to a fast-acting antihistamine, a peanut allergic who smells peanuts may potentially use transcranial stimulation to disrupt the engram. Furthermore, in the case of an allergic reaction triggered by peanut smells that activate an engram and not by direct exposure to peanut allergens, a transcranial magnetic stimulation that disrupts the corresponding neuronal network may be more efficacious than the administration of fast-acting antihistamines or injectable adrenaline because these do not target the pathogenic mechanism. Just as with the development of a theory of neuroimmune interactions, the investigation of such potential clinical interventions is a long-term endeavour, requiring the coordinated efforts of biologists and neuroscientists, mathematicians and computer scientists, psychologists and clinicians.

Interestingly, in the original 1886 article, Mackenzie reports putting an end to the pathogenic “engram” that was driving the ‘rose cold’ by having his patient examine the counterfeit flower, which significantly decreased subsequent reactions. Reflecting on insights on general psychosomatic phenomena emerging from this case, he concludes:

> *“If man’s relation to the external agents which are made to minister to his pleasure be thus perverted, is it not more rational to conclude that the disturbance arises from a defect in the subject himself, or some disturbance of the sentient apparatus, than from a subversion of the teleological purposes of natural objects?”*

Seen in a more modern content, this observation naturally suggests that empowering the patient by explaining the neuroimmunological mechanisms that underlie such engrams may (at least in the near term) be indeed the best approach for dealing with these neuro-pathogenic mechanisms of disease.

