## Supplementary Materials for "A Rose by any Other Name: Towards a quantitative theory of memory in neuroimmune memories" (from another version of this preprint: version 2, 2024-08-08)

### A Rose by any Other Name: Towards a quantitative theory of neuroimmune interactions Supplementary Information

#### A Mechanistic mathematical model derivation

The Engram-Immune Model (EIM), appearing in the body of the main paper, arises from the incorporation of various mechanistic processes in the immune system and the engram network. In this section, we provide a more detailed description of these mechanistic processes and the arising non-dimensionalisation.

The (dimensional) engram signal, $S$, is promoted in the presence of immune lesion and can self-promote via some promotion function $p(S)$. However, in the presence of the feedback signal $A$, $p(S)$ will decrease in strength and, above some 'engram threshold' $A^*$, will cause $S$ to downregulate its expression. Combining these elements provides the following ODE for the engram:

$$\frac{\mathrm{d}S}{\mathrm{d}t} = k_0(A^* - A)p(S) + k_1 V. \tag{A1}$$

To ensure a finite-time cut-off of the engram signal, we adopt the following modified-exponential growth rate

$$p(S) = \gamma_S S^\beta + S, \qquad 0 < \beta < 1, \tag{A2}$$

to ensure that $S$ can achieve a value of zero in finite time, unlike the exponential model. Nevertheless, at higher strengths of $S$, the dominant reaction kinetics return to the exponential model.

Next, to model the immune response $I$, we assume that the immune response is promoted in direct proportion to $S$ and that the feedback signal reduces $I$ down to some (positive) baseline level of immune activity, $I^*$. The immune response will naturally subside with decay rate $\delta_1$. In instances where the increase in immune response also damages the immune system itself, we incorporate a Michaelis-Menten-type damage response mechanism $\dfrac{SI}{S_0 + S}$. More general Hill-type response mechanisms are omitted due to emergence of bistability of non-zero equilibria and are viewed as nonphysical. Combining all of the aforementioned mechanisms provides

$$\frac{\mathrm{d}I}{\mathrm{d}t} = k_2 S - \frac{KSI}{S_0 + S} - \delta_1(I - I^*)(1 + \hat{K}A) + k_3 V. \tag{A3}$$

To model the immune lesion $V$, it is important to consider if the immune lesion is self-replicating or not. For allergic reactions, the immune lesion is introduced at some initial concentration $v_0$, but does not self-replicate.

However, for viral or bacterial infections, we can incorporate self-replication of the immune lesion using standard choices of growth mechanisms (e.g., logistic growth or otherwise, denoted as $f(V)$). To eliminate the immune lesion, the immune response reduces the lesion using the same modified-exponential function form as $p(S)$:

$$\frac{\mathrm{d}V}{\mathrm{d}t} = k_V f(V) - k_4 I(\gamma_V V^\beta + V). \tag{A4}$$

Finally, the feedback signal $A$ is created in the presence of an immune response above its baseline, including the aforementioned immune response that eliminates the lesion. Additionally, we will assume that the off-signal naturally decays from the system in the absence of patient insult:

$$\frac{\mathrm{d}A}{\mathrm{d}t} = Mk_4 I(\gamma_V V^\beta + V) + k_5(I - I^*) - \delta_2 A. \tag{A5}$$

These ODEs are coupled with initial conditions corresponding to baseline immune activity, inactive engram activity, and a non-zero immune lesion present:

$$(S(0), I(0), A(0), V(0)) = (0, I^*, 0, v_0). \tag{A6}$$

#### A.1 Non-dimensionalisation of the Engram-Immune Model

To produce a non-dimensional version of the EIM, we use the rescalings

$$S = \sigma x := \frac{k_0^2 A^{*2}}{k_2 k_4} x, \qquad I = I^* + \frac{k_0 A^*}{k_4} y, \qquad A = A^* z, \qquad V = \frac{M}{A^*} u, \qquad t = \frac{\hat{t}}{k_0 A^*}. \tag{A7}$$

By dropping hat notation for simplicity, we obtain the non-dimensional Engram-Immune Model:

$$\frac{\mathrm{d}x}{\mathrm{d}t} = (1 - z)(x + \zeta_1 x^\beta) + \Lambda_1 u, \tag{A8}$$

$$\frac{\mathrm{d}y}{\mathrm{d}t} = x - \frac{\kappa x(y + y_0)}{\alpha + x} - \delta y(1 + \lambda z) + \Lambda_2 u, \tag{A9}$$

$$\frac{\mathrm{d}z}{\mathrm{d}t} = (y + y_0)(u + \zeta_2 u^\beta) + \xi y - \eta z, \tag{A10}$$

$$\frac{\mathrm{d}u}{\mathrm{d}t} = -(y + y_0)(u + \zeta_2 u^\beta) + \Gamma f(u), \tag{A11}$$

where

$$\zeta_1 = \gamma_S \sigma^{\beta - 1}, \quad \zeta_2 = \frac{\gamma_V M^{\beta-1}}{A^{*\,\beta-1}}, \quad \delta = \frac{\delta_1}{k_0}, \quad \kappa = \frac{K}{k_0 A^*}, \quad \xi = \frac{k_5}{k_4 A^*},$$

$$\Lambda_1 = \frac{k_1 k_2 k_4}{k_0^3 A^{*\,2} M}, \quad \Lambda_2 = \frac{k_3 k_4}{k_0^2 A^* M}, \quad \lambda = \hat{K} A^*, \quad \eta = \frac{\delta_2}{k_0 A^*}, \quad \alpha = \frac{S_0}{\sigma}, \quad y_0 = \frac{I^* k_4}{k_0 A^*}.$$

Additionally, the initial condition of the EIM becomes

$$(x(0), y(0), z(0), u(0)) = (0, 0, 0, V_0), \quad \text{where} \quad V_0 = \frac{M v_0}{A^*}.$$
